# The role of Fnr paralogs for controlling anaerobic metabolism in diazotroph *Paenibacillus polymyxa* WLY78

**DOI:** 10.1101/2020.01.03.894683

**Authors:** Haowen Shi, Yongbin Li, Tianyi Hao, Xiaomeng Liu, Xiyun Zhao, Sanfeng Chen

## Abstract

Fnr is a transcriptional regulator that controls the expression of a variety of genes in response to oxygen limitation in bacteria. Genome sequencing revealed four genes (*fnr1*, *fnr3*, *fnr5* and *fnr7*) coding for Fnr proteins in *Paenibacillus polymyxa* WLY78. Fnr1 and Fnr3 showed more similarity to each other than to Fnr5 and Fnr7. Also, Fnr1 and Fnr3 exhibited high similarity with *Bacillus cereus* Fnr and *Bacillus subtilis* Fnr in sequence and structures. Deletion analysis showed that the four *fnr* genes, especially *fnr1* and *fnr3,* have significant impacts on the growth and nitrogenase activity. Single deletion of *fnr1* or *fnr3* led to 50% reduction in nitrogenase activity and double deletion of *fnr1* and *fnr3* resulted to 90% reduction in activity. Both of the aerobically purified His-tagged Fnr1 and His-tagged Fnr3 in *Escherichia coli* could bind to the specific DNA promoter. Genome-wide transcription analysis showed that Fnr1 and Fnr3 indirectly activated expression of *nif* (nitrogen fixation) genes and Fe transport genes under anaerobic condition. Fnr1 and Fnr3 inhibited expression of the genes involved in aerobic respiratory chain and activated expression of genes responsible for anaerobic electron acceptor genes.

**IMPORTANCE:** *Paenibacillus* is a genus of Gram-positive, facultative anaerobic and endospore-forming bacteria. The members of nitrogen-fixing *Paenibacillus* have great potential use as a bacterial fertilizer in agriculture. However, the functions of *fnr* gene(s) in nitrogen fixation and other metabolisms in *Paenibacillus* spp. are not known. Here, we revealed that copy numbers vary largely among different *Paenibacillus* species and strains. Deletion and complementation analysis demonstrated that *fnr1* and *fnr3* have significant impacts on the growth and nitrogenase activity. Both of the aerobically purified His-tagged Fnr1 and His-tagged Fnr3 purified in *Escherichia coli* could bind to the specific DNA promoter as *Bacillus cereus* Fnr did. Fnr1 and Fnr3 indirectly activated *nif* expression under anaerobic condition. Fnr1 and Fnr3 directly or indirectly activated or inhibited expression of many important genes involved in respiration, energy metabolism, Fe uptake and potentially specific electron transport for nitrogenase under anaerobic condition. This study not only reveals the roles of *fnr* genes in nitrogen fixation and anaerobic metabolism, but also provides insight into the evolution and regulatory mechanisms of *fnr* in *Paenibacillus*.

## INTRODUCTION

Most biological nitrogen fixation is catalyzed by molybdenum-dependent nitrogenase, which is distributed within bacteria and archaea. This enzyme is composed of two metalloproteins: MoFe protein and Fe protein (1). Nitrogenase is an oxygen-sensitive enzyme, and both the MoFe and Fe proteins are irreversibly damaged by oxygen (2). O_2_ exposure leads to inappropriate oxidation of the metalloclusters, decrease of protein secondary structure and further degradation (3). Exposure to oxygen irreversibly inactivates the Mo-, V-, and Fe-nitrogenases (3–5). To avoid oxygen inactivation, diazotrophs (nitrogen-fixing organisms) have evolved different strategies. One of the strategies is to tightly control the transcription of nitrogen fixation genes (*nif*) in response to the external oxygen concentration.

Fnr (fumarate and nitrate reduction) protein is a global regulator that binds a [4Fe–4S] cluster to monitor the oxygen status in the cell and then controls transcription of lot of genes in response to changes in oxygen levels (6–9). Fnr is widely distributed in Gram-negative bacteria (e.g. *Escherichia coli*) (10) and Gram-positive bacteria (e.g. *Bacillus subtilis*) (11). Fnr-related transcriptional regulators of the Crp/Fnr (cyclic AMP-binding protein/fumarate nitrate reduction regulatory protein) family have been reported to be involved in nitrogen fixation of some Gram-negative diazotrophs (12–15). For example, Fnr proteins are indirectly involved in controlling the activity of NifA in *Herbaspirillum seropedicae* SmR1 by regulating respiratory activity in relation to oxygen availability (16, 17). Fnr protein of *Klebsiella oxytoca* is required to relieve inhibition of NifA activity by its partner regulatory protein NifL under anaerobic conditions (14). In symbiotic *Bradyrhizobium japonicum* and *Sinorhizobium meliloti*, transcription of *nifA* and *fix* genes is predominantly controlled by the oxygen-responsive two component FixL–FixJ system, together with FixK which is a member of the Crp/Fnr superfamily, or by the redox-sensing system RegS–RegR (12, 13). In *Rhizobium leguminosarium* UPM791 FnrN is responsible for the expression of the high affinity oxidase encoded by *fixNOQP* which supports growth under microaerobic conditions and is essential for nitrogen fixation (15).

*Paenibacillus polymyxa* WLY78 can fix nitrogen under anaerobic or microaerobic and nitrogen-limited conditions and has a *nif* operon composed of 9 genes (*nifBHDKENXhesAnifV*) under the control of a σ^70^-dependent promoter in front of *nifB* gene (18). Recently, we have revealed that GlnR of *P. polymyxa* WLY78 activates *nif* transcription under anaerobic and nitrogen-limited condition, but GlnR together with glutamine synthetase (GS, *glnA* product) represses *nif* transcription under excess nitrogen and anaerobic condition (19).

Here, we searched the genome of *P*. *polymyxa* WLY78 and found that there are four genes coding for Fnr proteins. A total of 12 *fnr* deletion mutants, including single, double, triple and quadruple *fnr* deletion mutants were constructed by homologous recombination. The growth rates and nitrogenase activities among these *fnr* mutants and wild-type *P. polymyxa* WLY78 were comparatively analyzed. Each of the single deletion mutants Δ*fnr1*, Δ*fnr3*, Δ*fnr5* and Δ*fnr7* was effectively complemented by its corresponding *fnr* gene and by *B. subtilis fnr*. His-tagged Fnr1 and His-tagged Fur3 proteins expressed and purified in *E. coli* under aerobic conditions were used to verify the target genes by EMSA. Genome-wide transcription analysis in *P. polymyxa* WLY78 and the double mutant Δ*fnr13* were performed.

## RERULTS

### Identification of *fnr* genes in *P. polymyxa*

Analysis of the *P. polymyxa* WLY78 genome showed four *fnr*-like genes (named as *fnr1* (S6001676), *fnr3* (S6003218), *fnr5* (S6004820) and *fnr7* (S6005182)) (18). There are 39.98-53.63% identity among the four Fnr1, Fnr3, Fnr5 and Fnr7 proteins of *P. polymyxa* at amino acid level (Table S1). The highest (53.63%) identity was found between Fnr1 and Fnr3. Fnr1 and Fnr3 are more similar to each other than to Fnr5 and Fnr7. Like *P. polymyxa* WLY78, the three strains *P. polymyxa* M1, *P. polymyxa* E681, and *P. polymyxa* SC2 have four *fnr* genes. Each of the four *fnr* genes shows 99.44-100% identity with its corresponding gene from the different *P. polymyxa* strains (Table S1). However, some *Paenibacillus* species or strains, such as *Paenibacillus polymyxa* EBL06, *Paenibacillus polymyxa* Sb3-1 and *Paenibacillus jamilae* NS115 have only one Fnr which has 16.34-34.78% identity with the four Fnr proteins of *P. polymyxa* WLY78. Also, Fnr1, Fnr3, Fnr5 and Fnr7 proteins of *P. polymyxa* share 50.94%, 52.99%, 45.45% and 43.19% identities with of *B. subtilis* Fnr protein, respectively. Whereas, Fnr1 and Fnr3. Also, Fnr1, Fnr3, Fnr5 and Fnr7 proteins of *P. polymyxa* have 41.57%, 45.08%, 19.01% and 21.82% identities with *Bacillus cereus* Fnr.

The four Fnr proteins of *P. polymyxa* WLY78 contain the predicted N-terminal receiver domain and C-terminal DNA-binding domain (Fig. 1A), which represents the feature of the Crp/Fnr family protein (7). The [4Fe–4S] ^2+^ cluster of *B. subtilis* Fnr is coordinated by three C-terminally located cysteine residues at positions 227, 230, and 235 and one aspartate residue at position 141 (7, 20). Similar to *B. subtilis* Fnr, Fnr1 and Fnr3 proteins of *P. polymyxa* WLY78 have these conserved cysteine and aspartate residues. But Fnr5 and Fnr7 proteins of *P. polymyxa* WLY78 lack these conserved residues (Fig. 1A). The data suggest that Fnr1 and Fnr3 proteins of *P. polymyxa* WLY78 show high similarity with *B. subtilis* Fnr and *B. cereus* Fnr in sequence and structure.

**FIG 1.**
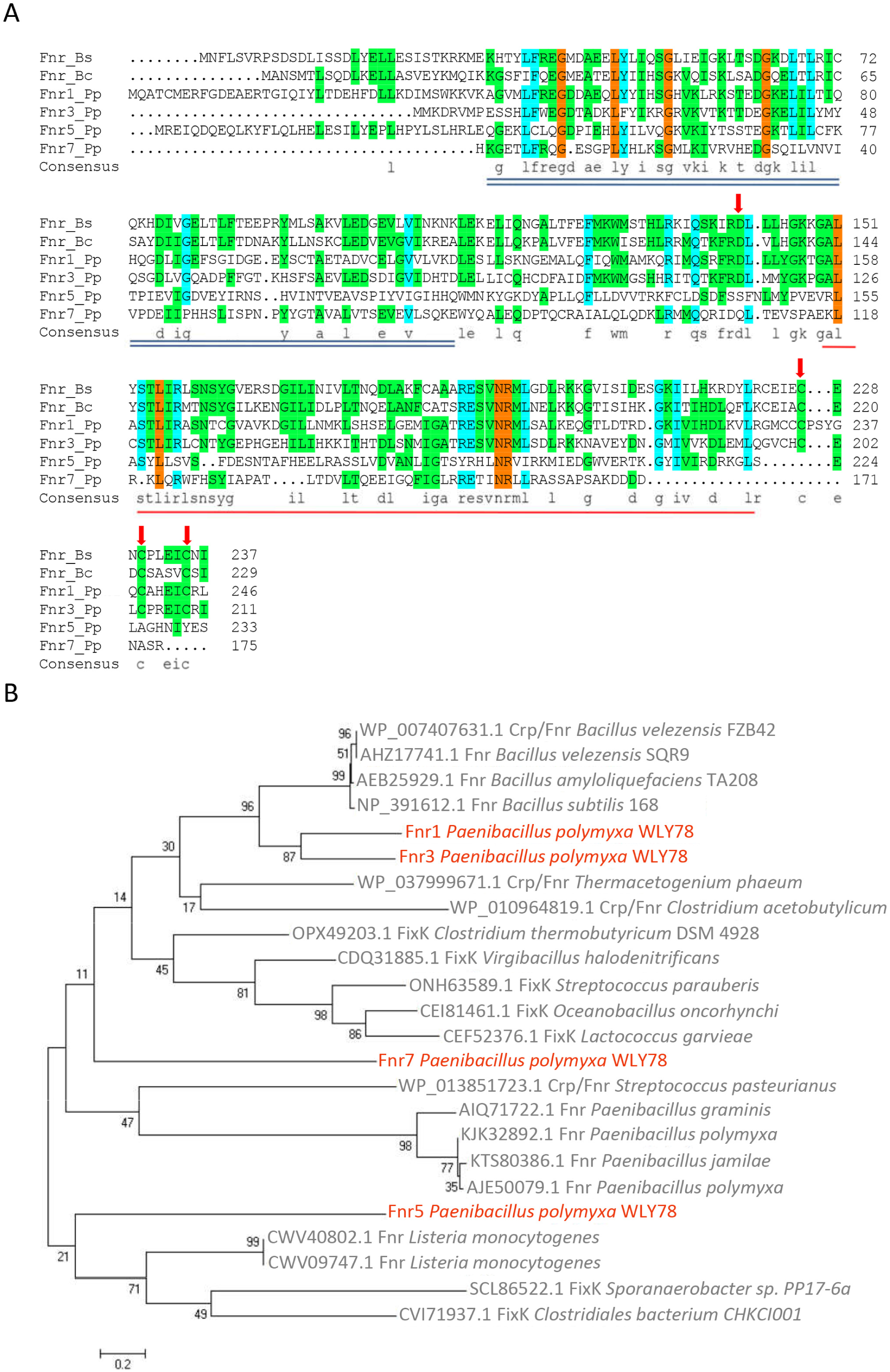
Homology analysis of Fnr proteins and phylogenetic analysis selected from Crp/Fnr superfamily in Firmicutes. (A) Alignments of Fnr proteins from *P. polymyxa*, *B. cereus* and *B. subtilis.* Conserved cysteines required for binding of the [4Fe-4S]^2+^ are indicated by red arrows. The double underlined sequence represents the region of the N-terminal DNA-binding domain. The red underlined sequence represents the region of sensory domain. Bs, *B. subtilis*; Bc, *B. cereus*; Pp, *P. polymyxa* WLY78 (B) The phylogenetic tree was constructed using neighbor joining method, with the bootstrap analyses of 1000 cycles.

A phylogenetic analysis showed that the four *P. polymyxa* Fnr proteins followed into 3 groups (Fig. 1B). Fnr1 and Fnr3 are in the clade with the Fnr group of *Bacillaceae*. Fnr5 is near the clade with the Fnr group from *Listeria* and FixK group of *Sporanaerobacter* and *Clostridiales*, while Fnr7 is divergent from Fnr and FixK group of *Bacillaceae*. The data are consistent with the protein homology analysis.

### Influence of *fnr* on growth under anaerobic condition

To explore the regulatory function of the four Fnr proteins of *P. polymyxa* WLY78, 12 unmarked *fnr* deletion (Δ*fnr*) mutants, including single, double, triple and quadruple deletion mutants, were constructed as described in the Methods. The number in the Δ*fnr* mutant indicates which *fnr* gene is deleted (e.g. Δ*fnr1* indicates deleting *fnr1* gene, Δ*fnr13* indicating deleting both *fnr1* and *fnr3* genes).

As Fnr protein is known to sense oxygen and plays a major role in altering gene expression during the switch from aerobic to oxygen limiting conditions, the influence of *fnr* on growth of *P. polymyxa* WLY78 under anaerobic conditions is here investigated. *P. polymyxa* WLY78 and multiple *fnr* deletion mutants were cultivated in nitrogen deficient medium with casamino acid under anaerobic and aerobic conditions (Fig. 2A). Except for the double *fnr* deletion mutant Δ*fnr57*, all of the *fnr* deletion mutants showed lower growth rate than *P. polymyxa* WLY78 did. Compared to wild-type *P. polymyxa* WLY78, each single *fnr* deletion mutant showed slow growth rate. The quadruple *fnr* deletion mutant Δ*fnr1357* showed the lowest growth rates among all of the 12 Δ*fnr* mutants, suggesting that the four *fnr* genes play roles under anaerobic condition. Notably, the single deletion mutants Δ*fnr1* and Δ*fnr3* and the double *fnr* deletion mutant Δ*fnr13* showed very low growth rate, suggesting that Fnr1 and Fnr3 proteins play an important role in anaerobic metabolisms in response to oxygen.

**FIG 2.**
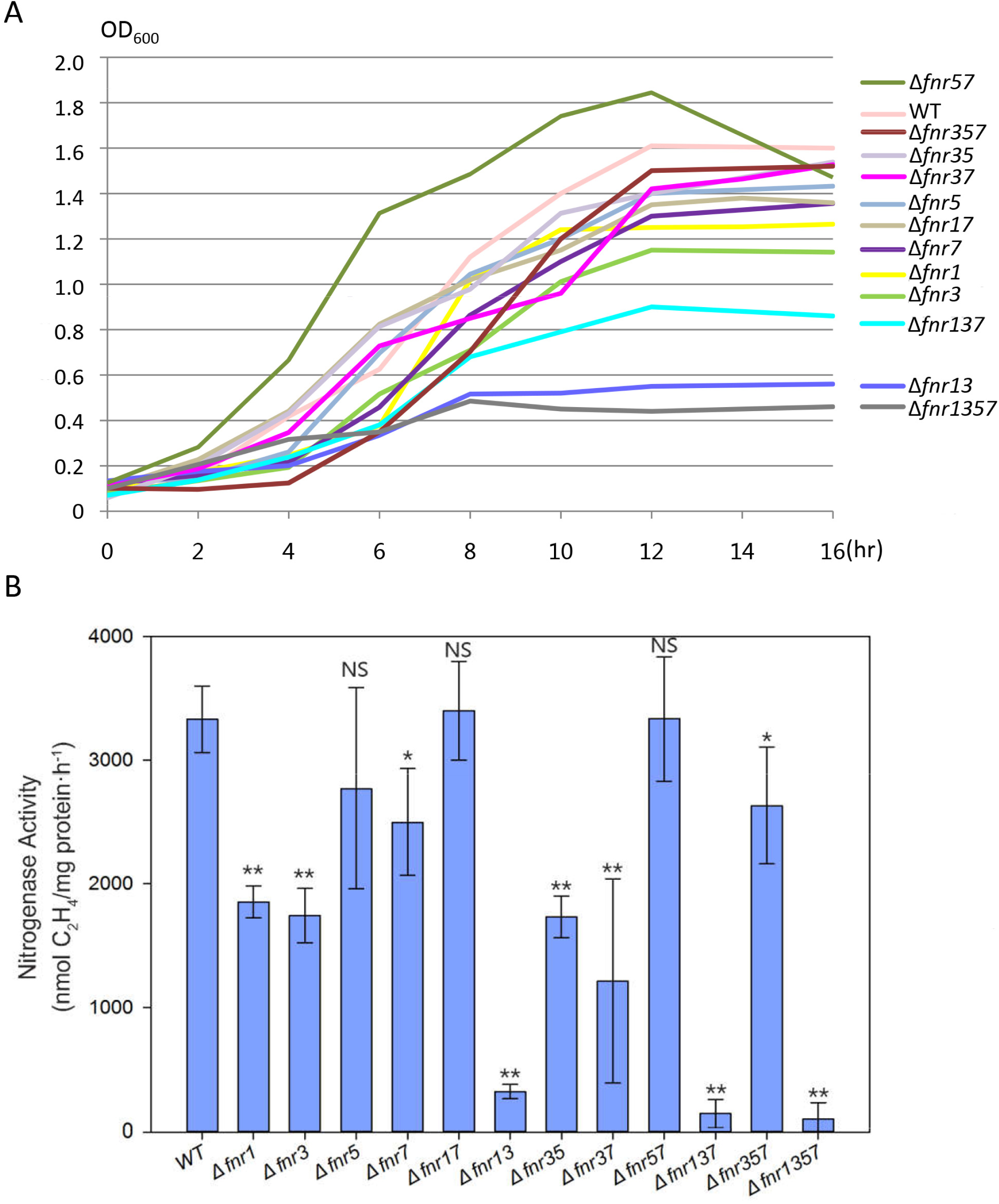
The growth curve and nitrogenase activity of the *fnr* deletion mutants. (A) Influence of the *fnr* deletion on growth under anaerobic condition. *P. polymyxa* WLY78 and the *fnr* deletion mutants were cultivated in nitrogen deficient medium with casamino acid and no oxygen. (B) Influence of the *fnr* deletion on nitrogenase activity under anaerobic condition. The nitrogenase activity of *P. polymyxa* WLY78 and the *fnr* deletion mutants was measured by acetylene reduction assay when grown anaerobically in nitrogen deficient medium.

### Effects of *fnr* on nitrogenase activity

Since nitrogenase is very sensitive to O_2_, nitrogen fixation is performed in anaerobic or microanaerobic conditions. To determine if Fnr proteins are related to nitrogen fixation, the nitrogenase activities of wide-type *P. polymyxa* WLY78 and multiple *fnr* deletion mutants grown anaerobically in nitrogen deficient medium were measured by using the method of the reduction of acetylene to ethylene (21, 22). As shown in Fig. 2B, the nitrogenase activities of Δ*fnr1* and Δ*fnr3* were decreased to about 50% of the wild type, while the activities of Δ*fnr5* and Δ*fnr7* were decreased to about 73-79% of the wild type. And the nitrogenase activity of Δ*fnr37* and Δ*fnr13* were decreased to about 36% and 10% of the wild type, respectively. Notably, the nitrogenase activities of Δ*fnr137* and Δ*fnr1357* were nearly lost. The data are consistent with the growth rates of these mutants observed as above. The results imply that the four *fnr* genes, especially *fnr1* and *fnr3,* play roles in nitrogen fixation.

Furthermore, complementation of Δ*fnr1,* Δ*fnr3,* Δ*fnr5* and Δ*fnr7* with its corresponding *P. polymyxa fnr* gene and *B. subtilis fnr* gene under the control of its own promoter was performed. As shown in Fig. S1, *fnr1, fnr3, fnr5* and *fnr7* from *P. polymyxa* WLY78 in complemented strains (Δ*fnr1C*, Δ*fnr3C*, Δ*fnr5C* and Δ*fnr7C*) restored the nitrogenase activity of its corresponding mutant to more than 90% activity of wild type. Complementation with His-tagged Fnr1 and His-tagged Fnr3 in complemented strains (Δ*fnr1Chis* and Δ*fnr3CHis*) also restored the nitrogenase activity of its corresponding mutant to more than 90% activity of the wild type. Moreover, we found that *B. subtilis fnr* gene greatly improved the nitrogenase activity of the four single *fnr* deletion mutants, especially the activities of Δ*fnr1* and Δ*fnr3*. Also, *B. subtilis fnr* gene greatly restored the nitrogenase activities of the multiple deletion mutants Δ*fnr13,* Δ*fnr137* and Δ*fnr1357.* The data confirm that the four *fnr* genes of *P. polymyxa* WLY78, especially *fnr1* and *fnr3*, play an important role in nitrogen fixation. The data suggest that Fnr1 and Fnr3 of *P. polymyxa* and Fnr of *B. subtilis* are similar in function.

### The *nifH* transcription in the *fnr* deletion mutants

The *nifH* transcriptions in different mutants were assayed by qRT-PCR (Fig. S2A). The *nifH* in Δ*fnr3* was expressed at basic level. The *nifH* transcriptions in Δ*fnr1*, Δ*fnr5* and Δ*fnr7* were decreased to about 40%, 70% and 90% of wild type, respectively. Whereas, the *nifH* transcriptions in Δ*fnr13* and Δ*fnr1357* were nearly lost. Furthermore, the effects of *fnr* on *nif* expression were performed by measuring the β-galactosidase activity of *P. polymyxa* WLY78 and *fnr* deletion mutants that carrying a transcriptional *lacZ* fusion to *nif* promoter region (P*nif-lacZ* fusion). Compared to wild type, mutants Δ*fnr1,* Δ*fnr3*, Δ*fnr13* and Δ*fnr1357* nearly lost β-galactosidase activities (Fig. S2B), in agreement with the nitrogenase activities and the *nifH* transcriptions in these *fnr* mutants.

### Prediction and verification of Fnr target genes

To decipher the Fnr regulon of *P. polymyxa* WLY78, its target genes were predicted. According to the known Fnr-binding sequence of *Bacillus* and *Paenibacillus* in RegPrecise (http://regprecise.lbl.gov), the PWM (position weight matrix) of Fnr-binding site was constructed using MEME (http://meme-suite.org). The Fnr-binding consensus motif composed of a 16 bp palindromic sequence 5’-TGTGA-N6-TCACA-3’ was determined. Then we used the 16 bp Fnr consensus binding-motif to scan the regions from -350 to +50 bp relative to the translational start codon (ATG) of genes in *P. polymyxa* WLY78 genome with the MAST application (http://meme-suite.org) (23). A total of 143 putative Fnr target genes with the E-value ≤ 10 (the smaller the E-value, the greater the probability) form *P. polymyxa* WLY78 genome were identified (Table 1). As annotated by the COG (Cluster of Orthologous Group), the 143 putative target genes were allocated to 12 groups by biological function. Of the 143 putative target genes, 19 belong to regulatory genes, 23 genes are related to energy metabolism, 11 genes are related to carbon metabolism, 54 genes are related to other metabolisms and 36 are the genes whose functions are unknown or unclassified (Table 1). As shown in Table 1, there is one Fnr-binding site in most of the 143 putative target genes, such as *fnr1*, *nark*, *narG,* and *resD.* There are two Fnr-binding sites in the promoter regions of the 16 genes, including *fnr3*, *fur3*, *adhC*, *adhE*, *adhP*, *cah*, *hmp*, *cydA*, *ndh*, *nemA*, *yugK*, *lacI*, *padR*, *accB*, *yphA* and *yhcN*. There are three Fnr-binding sites in the promoter regions of *nox* and *pflB*.

**Table 1.**
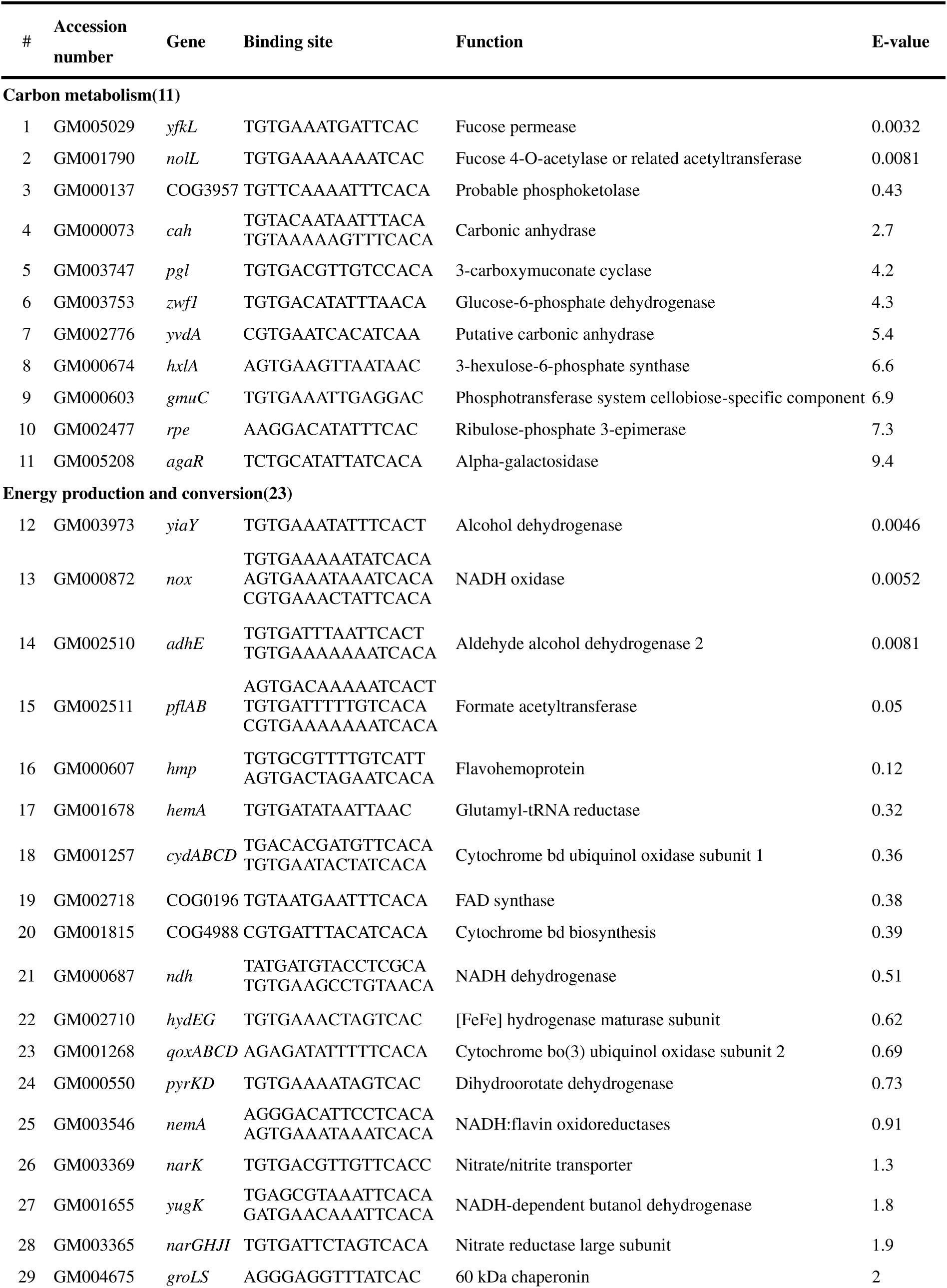
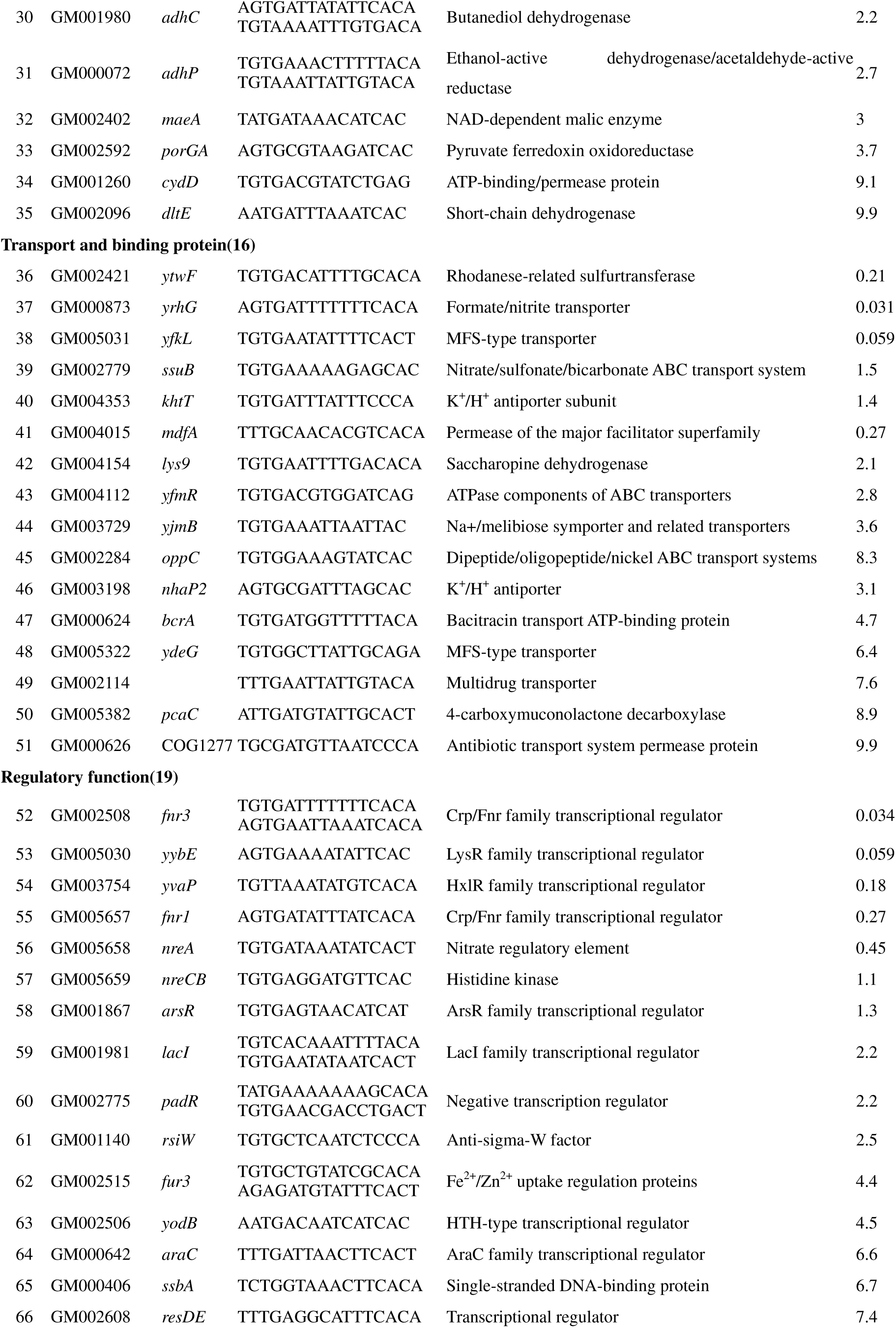
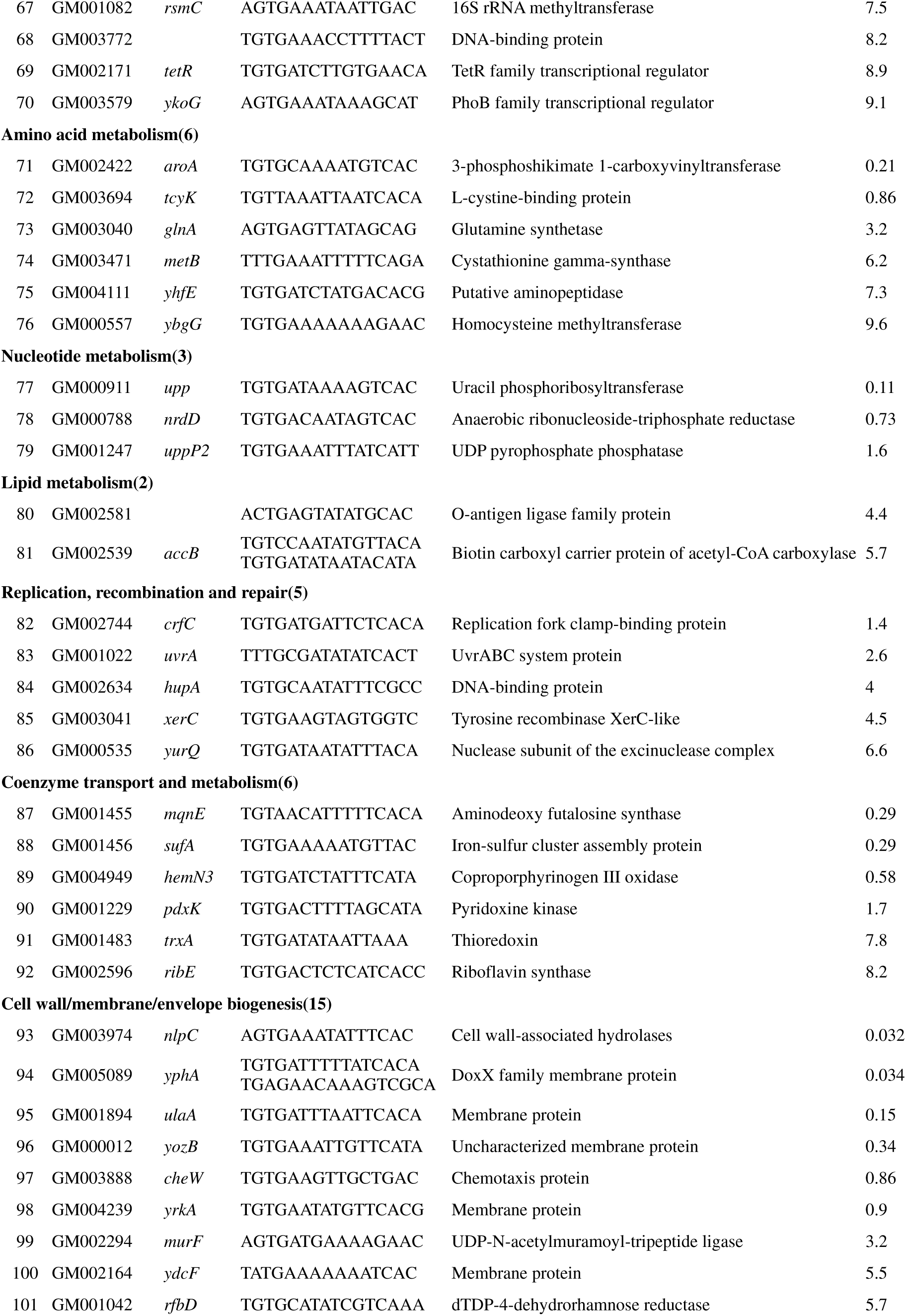
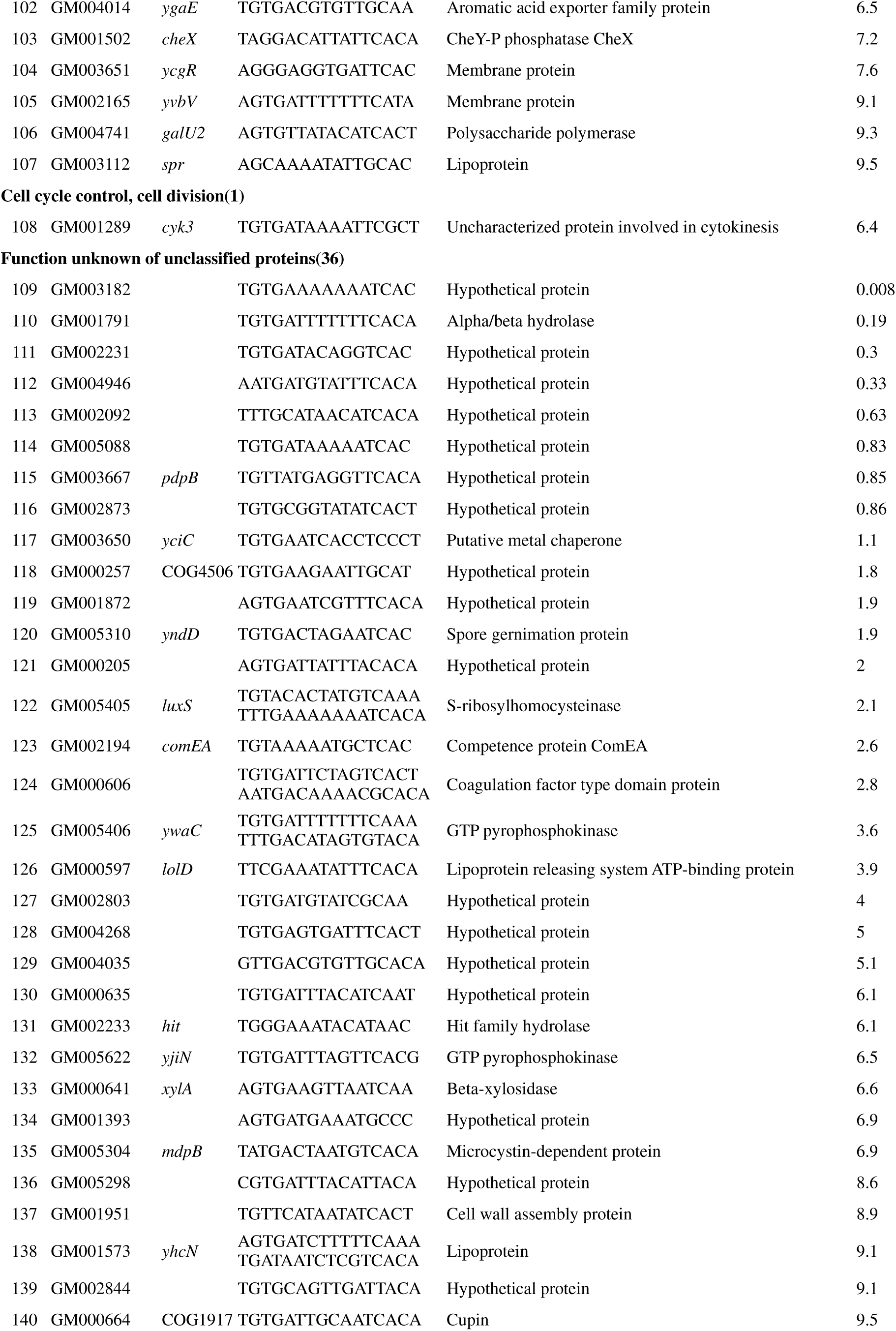
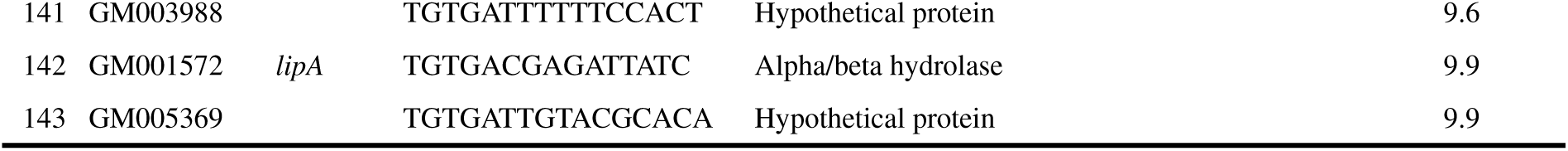
The prediction of Fnr target genes in *P. polymyxa* WLY78

The Fnr-binding motif was shown in Fig. 3A. To determine the accuracy of the bioinformatic analysis, the 13 promoter regions, including 11 putative targets with Fnr-binding sites and 2 target genes without predicted Fnr-binding sites were chosen to do electrophoretic mobility shift assays (EMAS). Fnr1 protein tagged with 6 Histidine at N-terminus (designated as NHis_6_-Fnr1) and Fnr3 protein tagged with 6 Histidine at C-terminus (designated as Fnr3-CHis_6_) were expressed and purified in *E. coli* under aerobic condition and then the two recombinant Fur1 and Fnr3 were used in EMSA. EMSA under aerobic condition showed that both Fnr1 and Fnr3 proteins could bind to the promoter regions with the predicted Fnr-binding sites of the 8 operons: *qoxABCD* (encoding cytochrome *aa3* quinol oxidase)*, narGHJI* (encoding nitrate reductase)*, ndh* (encoding NADH dehydrogenase)*, hemN3* (encoding oxygen-independent coproporphyrinogen-III oxidase)*, hydEG* (encoding [FeFe] hydrogenase)*, nrdDG* (encoding anaerobic ribonucleoside triphosphate reductase)*, pflBA* (encoding formate acetyltransferase)*, resDE* (encoding two-component regulatory proteins) (Fig. 3B). However, the promoter regions of *cydABCD* operon (encoding cytochrome *bd* ubiquinol oxidase) and *narK* (encoding nitrate/nitrite transporter) were only bound by Fnr3. Moreover, EMSA showed that neither Fnr1 nor Fnr3 could bind to the promoter regions of *nif* and *feoAB* (encoding the ferrous-iron transporter FeoAB), consistent with the facts that there was no Fnr-binding site in the promoter regions of these genes. However, EMSA showed that no binding of Fnr1 and Fnr3 to the promoter region of *glnRA* with Fnr-binding site.

**FIG 3.**
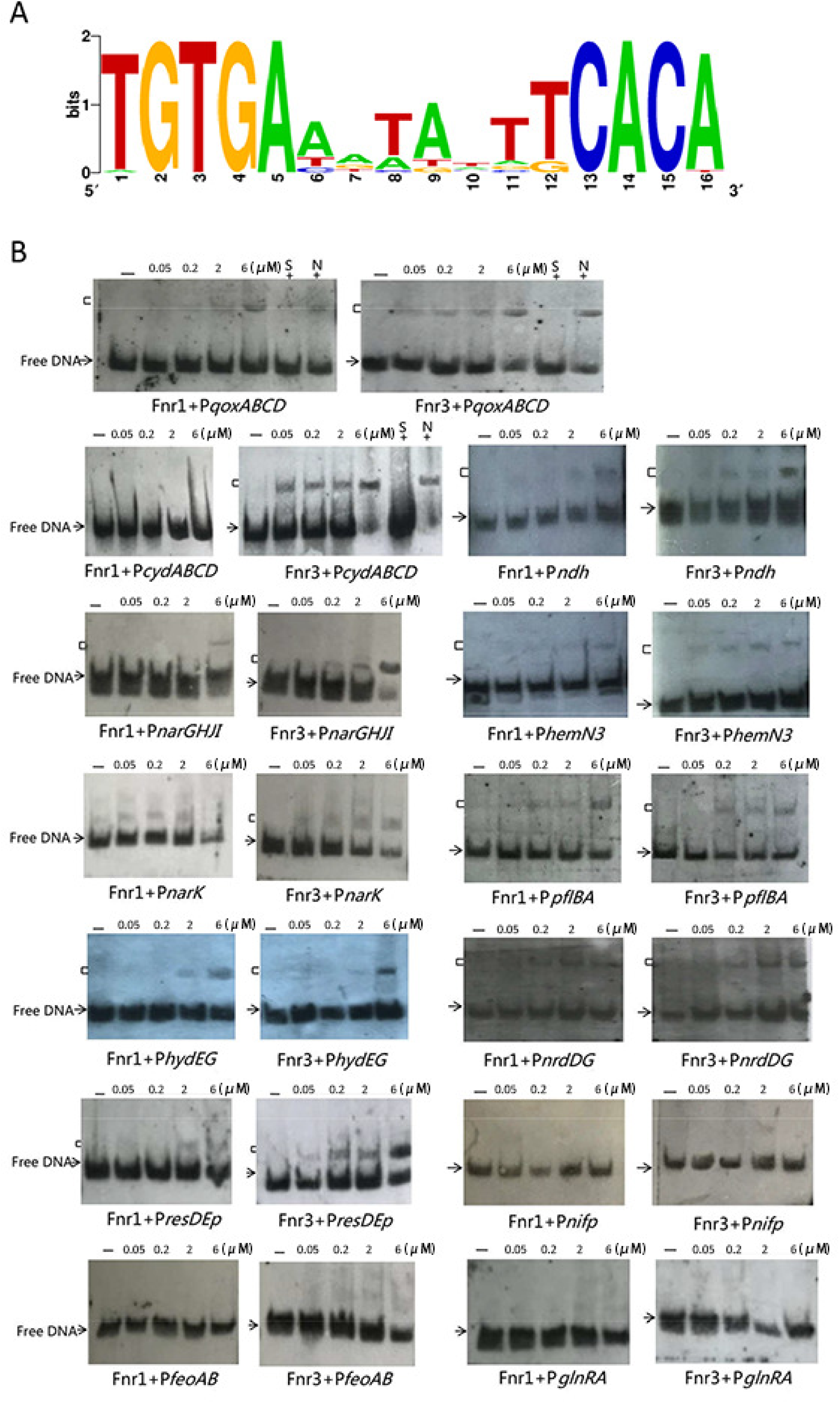
Fnr-binding sites predicted by software and verification by electrophoretic mobility shift assays (EMSA). (A) Consensus sequence of the predicted Fnr-binding sites. (B) *In vitro* binding of NHis_6_-Fnr1 and Fnr3-CHis_6_ to promoter region of some Fnr target genes. 179-442 bp DNA with final concentration of 0.03 pmol was used. The ‘-’ in Lane 1 indicates EMSAs without NHis_6_-Fnr1 or Fnr3-CHis_6_. Lanes 2–5 contained increasing concentrations (0.05, 0.2, 2, 6 μM) of NHis_6_-Fnr1 or Fnr3-CHis_6_ as indicated by the height of the triangle above the gel. S and N indicate competition assays with a 100-fold excess of unlabelled specific probe and nonspecific competitor DNA, respectively. Arrowheads: free probes. Brackets: DNA-protein complexes.

### RNA-Seq transcriptome analysis of wild-type and Δ*fnr13* strains

To assess the effects of Fnr1 and Fnr3 proteins on global gene expression in anaerobic condition, the genome-wide transcription analysis of *P. polymyxa* WLY78 and Δ*fnr13* mutant cultured under N_2_-fixing condition (without O_2_ and NH_4_^+^) was performed. Transcripts showed statistically significant differences with q-value (p-adjusted) ≤ 0.05 and a |log_2_ FC| ≥ 1 were accepted as candidate differential expression genes (DEGs). Of the 5661 genes contained in the genome of *P. polymyxa* WLY78, 301 genes, including 202 genes and operon, were differentially expressed in Δ*fnr13* compared to wild type (Table S2). Of the 301 genes, 116 were markedly up-regulated, indicating that they are directly or indirectly repressed by Fnr, and 185 were significantly decreased, suggesting that they were directly or indirectly activated by Fnr.

### Influence of *fnr* genes on transcription of the *nif* and *glnRA* genes

The 9 genes (*nifBHDKENXhesAnifV*) are organized as a *nif* operon in *P. polymyxa* WLY78. In this study, we find that the expression levels of the 9 genes within the *nif* operon in Δ*fnr13* were significantly down-regulated by 6.51-7.47 Log_2_FC (Fig. 4A). The data are consistent with the decreased nitrogenase activity and *nifH* transcription of Δ*fnr13* mutant. However, there was no predicted Fnr-binding site in the promoter region of the *nif* operon and EMSA also showed that Fnr1 or Fnr3 did not bind to the promoter region of the *nif* gene (Fig. 3B). These results indicated that Fnr1 and Fnr3 indirectly activated the expression of *nif* gene operon under anaerobic conditions. Expression of *glnRglnA* operon that plays regulatory role in *nif* transcription was up-regulated 1.71-1.74 log_2_FC.

**FIG 4.**
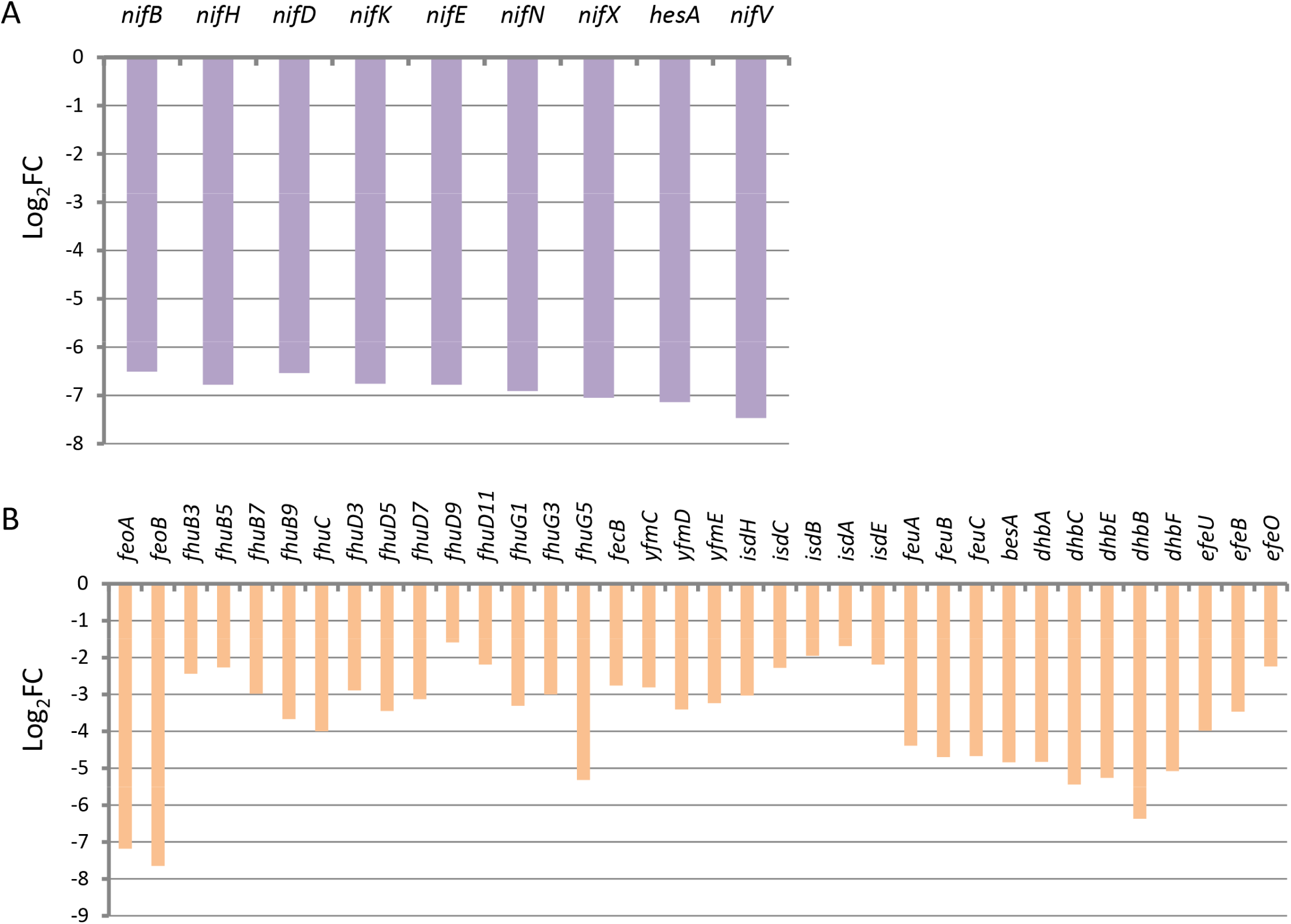
Differential expression of the *nif* genes and iron transport genes. (A) Differential expression of the 9 *nif* genes. (B) Differential expression of the genes involved in iron transport. FC in Log_2_FC indicates fold change (the read count ratio of Δ*fnr13* and wild type).

### Influence of the *fnr1* and *fnr3* genes on transcription of the Fe transporter genes

Fe is an essential element for nitrogenase. Fe is the soluble Fe^2+^ form (ferrous iron) under anaerobic condition or at acidic pH and the major route for bacterial ferrous iron uptake was via Feo (Ferrous iron transport) system composed of FeoA, FeoB and FeoC (24, 25). Fe at neutral pH is the poor solubility form of Fe^3+^ (ferric iron) which is often biologically unavailable (26). Many bacteria excrete ferric chelators, called siderophores, to take up Fe^3+^. Usually, bacteria take up ferric complexes, including ferric hydroxamate (FhuCDBA), ferric citrate (YfmCDEF), ferric-haem, ferric-bacillibactin uptake system (FeuABC) (27).

Our study showed that 36 Fe transporter genes in Δ*fnr13* mutant were down-regulated 1.59-7.65 Log_2_FC (Fig. 4B, Table S2). Of the 36 Fe transporter genes, only *feoAfeoB* operon is involved in Fe^2+^ uptake and the other 34 genes belong to Fe^3+^ transport systems. The highest differentially expressed genes *feoA* and *feoB* were down-regulated 7.18-7.65 Log_2_FC. The 13 *fhu* genes belonging to ferric hydroxamate system were down-regulated from 5.32 to 1.59 Log_2_FC. Especially, transcriptions of *yfmCDE* involved in ferric citrate transport system and *isdHCBAE* involved in ferric-haem transport system were also down-regulated in Δ*fnr13* mutant. The data are consistent with our recent reports that all of the Fe transporter genes were up-regulated in N_2_-fixing condition (without O_2_ and NH_4_ ^+^) (28). As described above, there are no Fnr-binding sites in the promoter regions of the 36 Fe transporter genes and EMSA also showed that Fnr1 or Fnr3 did not bind to the promoter region of the *feoAB* operon. Thus we deduce that Fnr1 and Fnr3 indirectly activated the expression of Fe transporter genes under anaerobic condition.

### Influence of *fnr* genes on transcription of respiration and energy metabolism genes

Based on the genome sequence, the respiratory chain of *P. polymyxa* WLY78 was shown in Fig. 5A. It is composed of several dehydrogenases that transfer electrons to an intramembrane pool of menaquinone and some terminal oxidases responsible for reoxidation of menaquinol. The terminal oxidases include at least two types: one consisting of a cytochrome *bd*-type quinol oxidase and the second one consisting of cytochrome *aa*3 oxidase.

**FIG 5.**
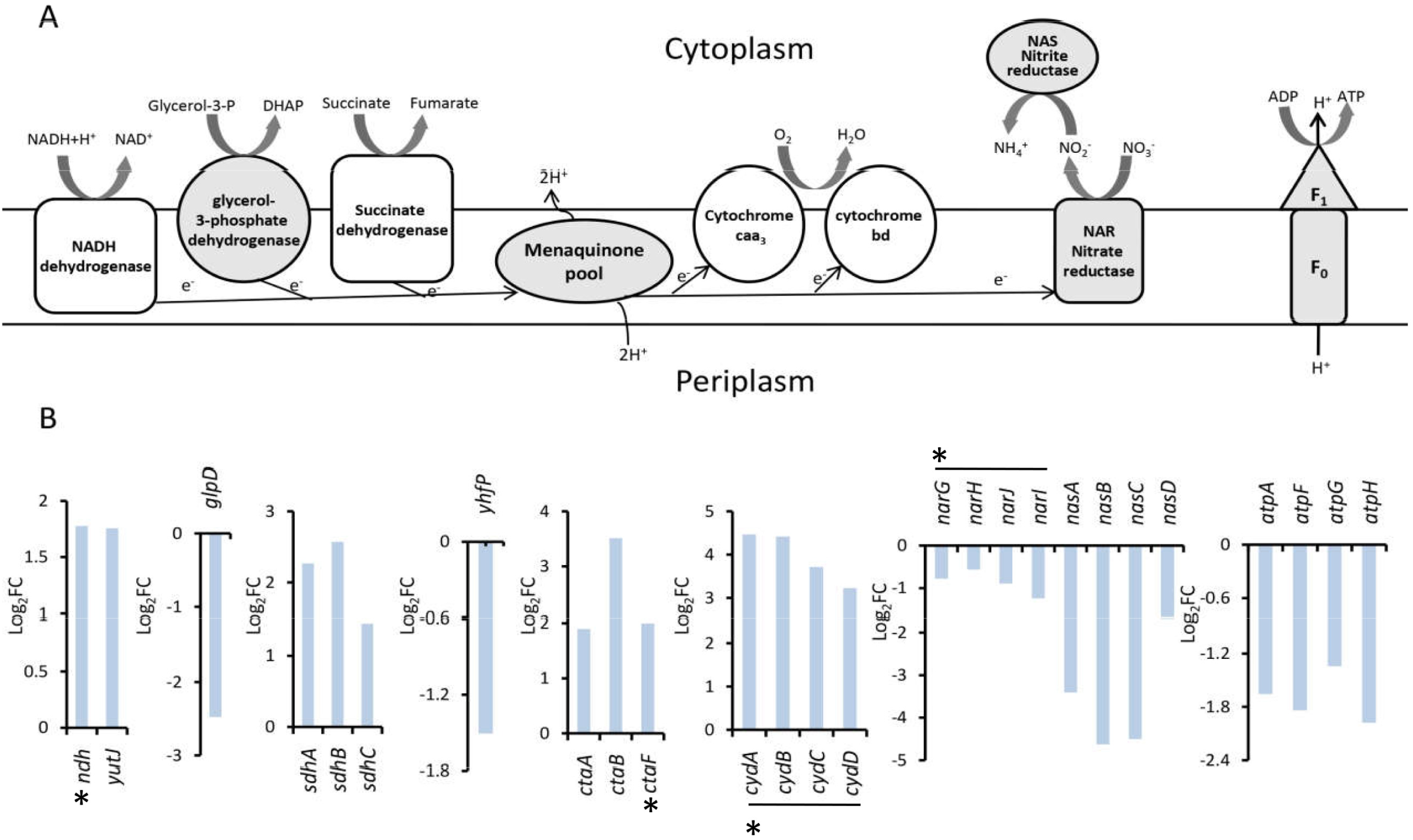
Differential expression of the genes related to anaerobic respiration and energy metabolism. (A) Schematic representation of the probable components of anaerobic respiration and energy metabolism in *P. polymyxa* WLY78 based on the genome annotation. Gray area represents the components whose transcripts are down-regulated in Δ*fnr13*. (B) Differential expression of the genes represented in the schema (A), FC in Log_2_FC indicates fold change. The horizontal line above or down genes indicates these genes are in the same transcription unit. The asterisk indicates that the promoter region of the gene contains predictive Fnr-binding site.

The dehydrogenases that play an important role in respiration in Gram-positive *Corynebacterium glutamicum* include a non-proton-pumping NADH dehydrogenase encoded by the *ndh* gene, malate:quinone oxidoreductase encoded by the *mqo* gene, and succinate dehydrogenase encoded by the *sdhCAB* genes (29). Here, we found that there were 13 genes encoding dehydrogenase were differentially expressed (Table S2). Of these genes, *ndh* and *sdhABC,* the major dehydrogenase genes in the respiratory chain were up-regulated 1.45-2.57 Log_2_FC. Other dehydrogenase genes, such as *yutJ* (NADH dehydrogenase), *yugK (*Probable NADH-dependent butanol dehydrogenase*), hcaD (*NAD(FAD)-dependent dehydrogenases*), ldh* (L-lactate dehydrogenase), were up-regulated 1.76-8.44 Log_2_FC, while *glpD*, *alkH* (aldehyde dehydrogenase), *adhE*, *fdhD* (formate dehydrogenase) and *adhP* genes were obviously down-regulated 2.48-5.87 Log_2_FC (Table S2). EMSA showed that NHis_6_-Fnr1 and Fnr3-CHis_6_ could bind to the promoter regions of *ndh* with the predicted Fnr-binding site (Fig. 3B). The *qoxABCD* encoding (cytochrome aa3-type oxidase) and cydABCD (encoding cytochrome bd-type oxidase) were up-regulated 1.8 to 4.5 Log_2_FC.

Many bacteria are able to grow anaerobically using alternative electron acceptors, including nitrate or fumarate (30). We found that anaerobic electron acceptor genes *narGHJI* (nitrate reductase, Nar), *nasABCD* (nitrite reductase, Nas) and *narK* (nitrate/nitrite transporter, NarK) were down-regulated from 0.6 to 4.6 fold in Δ*fnr13* mutant. As described above, the Fnr-binding sites in the upstream region of *narGHJI* and *narK* were predicted and confirmed by EMSA. Thus, the results suggest that Fnr1 and Fnr3 directly activate expression of *narGHJI* and *narK* in anaerobiosis and indirectly activate expression of *nasABCD*. The results are consistent with previous studies that the expression of *narGHJI* was intensely induced by anaerobic condition and the induction was dependent on Fnr in *B. subtilis* (31). In addition, the *atpAFGH* genes encoding ATP synthase were also down-regulated, but no Fnr-binding site was predicted in upstream regions of these genes, suggesting that Fnr1 and Fnr3 might indirectly activate expression of *atpAFGH* genes under anaerobic condition. These results indicated that Fnr1 and Fnr3 inhibited expression of genes involved in aerobic respiration process and activate express of genes involved in anaerobic energy metabolism. ResD-ResE (two-component regulatory proteins) and FNR were previously shown to be indispensable for nitrate respiration in *B. subtilis* (32, 33). Here we show that expression of *resDE* inΔ*fnr13* mutant were down-regulated 1.4-1.6 Log_2_FC, in agreement with our previous report that *resD* and *resE* were obviously up-regulated in *P. polymyxa*WLY78 under N_2_-fixation condition (without NH_4_^+^ and O_2_) (28). EMSA Fnr1 and Fnr3 could bind to the promoter of *resDE* operon with a Fnr-binding site, consistent with the report that *B. cereus* Fnr regulated expression of *resDE* (38). The data indicated that Fnr1 and Fnr3 inhibited expression of the genes involved in aerobic respiratory chain and activated expression of genes responsible for anaerobic electron acceptor genes

### Transcriptional analysis of the potential electron transporters for nitrogenase

Nitrogen fixation is carried out by the enzyme nitrogenase, which transfers electrons originating from low potential electron carriers, such as flavodoxin or ferredoxin molecules, to molecular N_2_ (34). A flavodoxin (encoded by *nifF*) mediates electron transfer from a pyruvate: flavodoxin oxidoreductase (encoded by *nifJ*) to the Fe protein of nitrogenase in *K. oxytoca* (34).

At present, we do not know how many genes and which gene are involved in electron transfer for nitrogenase in *P. polymyxa* WLY78. The differentially expressed genes that may be the potential electron transfer for nitrogenase in *P. polymyxa* WLY78 were shown in Table S2 and Fig. 6A. Homology analysis showed that *fldA* (encoding flavodoxin) of *P. polymyxa* WLY78 showed 30% identity with *K. oxytoca nifF*. Expression of *fldA* in Δ*fnr13* was down-regulated by 2.74 Log_2_FC. As shown in Fig. 6A, two transcripts *hydEG* located in plus strand and COG0196 *fdhF hycB hydAN aspA hydFG* located in minus strand were significantly down-regulated 7.13-12.55 Log_2_FC. *hydA* encodes Fe-Fe hydrogenase whose synthesis relies on maturation factors HydF (GTPase), HydE and HydG (35), while *hydB* encodes ferredoxin and *fdhF* encodes formate dehydrogenase. Each of the promoter regions of the two operons *hydEG* and COG0196 *fdhF hycB hydAN aspA hydFG* contain a predicted Fnr binding site, and EMSA showed that NHis_6_-Fnr1 and Fnr3-CHis_6_ could bind to the promoter of *hydEG* (Fig. 3B). It has been reported in *Clostridium*, electrons produced by the oxidation of pyruvate are transferred to the acceptor ferredoxin, and then the ferredoxin can act as electron donors to reduce Fe-Fe hydrogenase HydA to produce hydrogen (36). In addition, the expression of *hemN1*, *hemN3* (*hemN* encoded oxygen-independent coproporphyrinogen-III oxidase) and COG1249 (encoded FAD-dependent oxidoreductase) were also down-regulated 2.40-9.61 Log_2_FC. EMSA showed that Fnr1 and Fnr3 could bind to the promoter of *hemN3* with a predicted Fnr binding site. Furthermore, we found that *hmp* (flavohemoprotein), *wrbA* (multimeric flavodoxin), *ywnB* (NADH-flavin reductase), *ribE* (riboflavin synthase) and *groSgroL* (chaperonin) were up-regulated 1.26-6.88 Log_2_FC. EMSA showed that Fnr1 and Fnr3 could bind to the promoter of *hmp* with two Fnr binding sites. We deduce that some of the differentially expressed genes, including *fldA* (flavodoxin and *hydB* (ferredoxin) may be involved in be the potential electron transfer for nitrogenase in *P. polymyxa* WLY78. However, some genes involved in electron transfer for nitrogenase, such as *nfrA* (encoding NAD(P)H Flavin oxidoreductase), were not differentially expressed.

**FIG 6.**
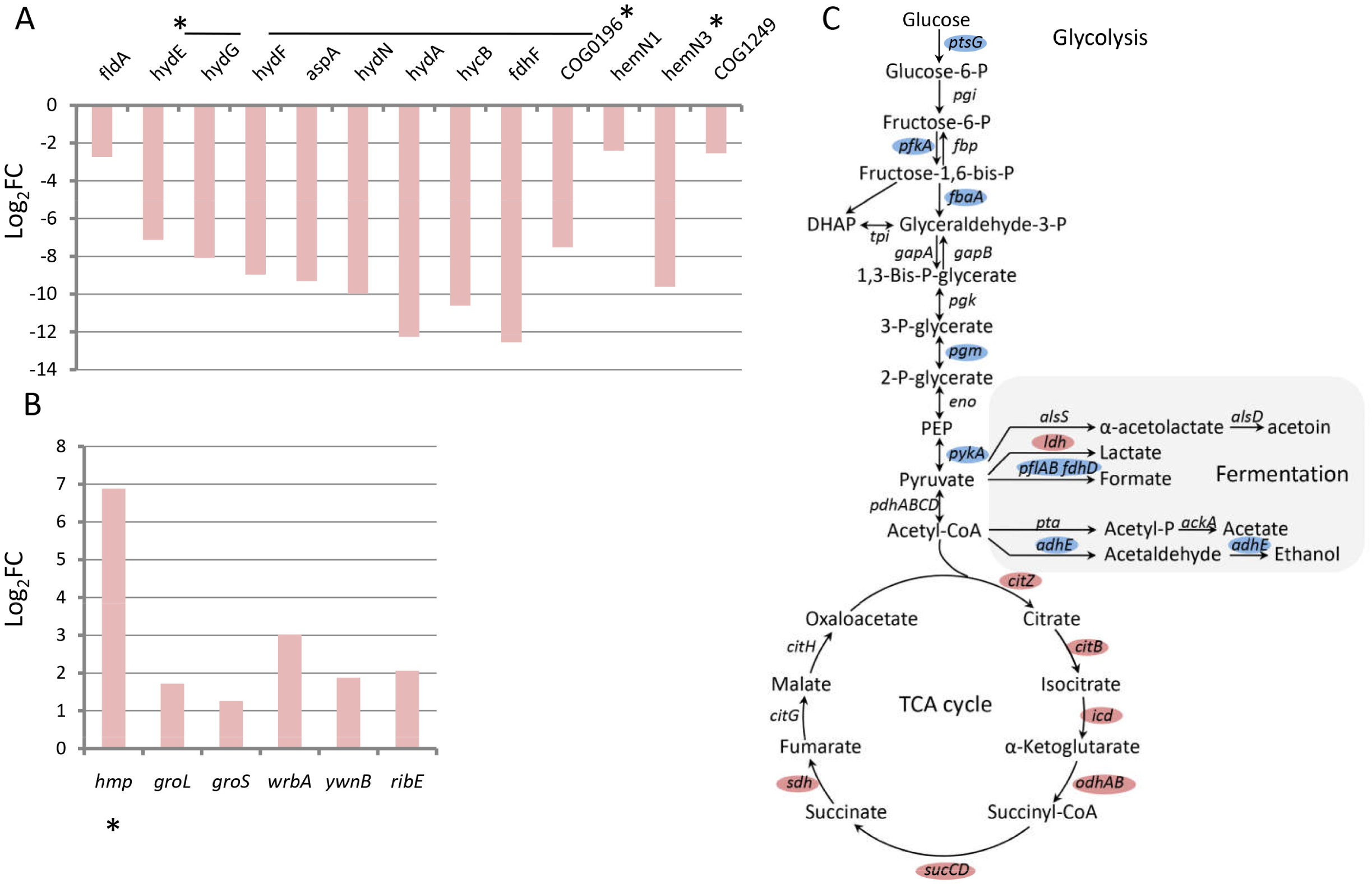
Differential expression of the genes related to electron transport and carbon metabolism. (A) Differential expression of the genes in electron transport. (B) Differential expression of the genes in carbon metabolism. (C) Schematic representation of the probable components of carbon metabolism (glycolysis, TCA cycle and fermentation) in *P. polymyxa* WLY78 based on the genome annotation. Blue and red indicate the components whose transcripts are down-regulated and up-regulated in Δ*fnr13*, respectively. For abbreviation: P, phosphate; DHAP, dihydroxyacetone phosphate; PEP, phosphoenolpyruvate. The horizontal line above genes indicates these genes are in the same transcription unit. Asterisk indicates that the promoter region of the gene contains the predictive Fnr-binding sites.

### Influence of *fnr* genes on transcription of genes involved in carbon metabolism

Genes involved in carbon metabolism (such as glycolysis, the Krebs (TCA) cycle and fermentation) were shown in Fig. 6C and Table S2. The down-regulated (1.33-3.28 Log_2_FC) genes in Δ*fnr13* were involved in glycolysis. These genes included *ptsG* (encoding glucose-specific component in PTS system), *pfkA* (encoding ATP-dependent 6-phosphofructokinase), *fbaA* (encoding fructose-bisphosphate aldolase), *pgm* (encoding β-phosphoglucomutase) and *pykA* (encoding pyruvate kinase). However, Fnr-binding sites were not found in the upstream regions of these genes, suggesting that Fur indirectly affected expression of the genes involved in glycolysis.

Many genes participated in formate and ethanol metabolism were significantly down-regulated in Δ*fnr13* mutant, such as *pflBA* (encoding formate acetyl transferase), *fdhD* (encoding formate dehydrogenase), *adhE* (encoding aldehyde-alcohol dehydrogenase) and *alkH* (encoding aldehyde dehydrogenase). Multiple Fnr-binding sites in the upstream regions of *pflBA* and *adhE* were predicted and EMAS also showed the binding of Fnr1 and Fnr3 to the promoter of *pflBA* (Table1 and Fig. 3B). This implied that Fnr1 and Fnr3 might have a direct regulation in expression of these genes under anaerobic condition. In contrast, *ldh* encoding L-lactate dehydrogenase was significantly up-regulated by 6.1 fold, but there was no predicted Fnr-binding site in the promoter region of this gene.

Many genes in the Krebs cycle were significantly up-regulated from 1.21 fold to 4.21 fold, and they included *citZ* (encoding citrate synthase), *citB* (encoding aconitate hydratase), *icd* (encoding isocitrate dehydrogenase), *odhAB* (encoding 2-oxoglutarate dehydrogenase), *sucCD* (encoding succinyl-CoA synthase) and *sdhABC* (encoding succinate dehydrogenase). However, there were no predicted Fnr-binding sites in the upstream region of these genes. These data suggested that Fnr1 and Fnr3 indirectly activated expression of genes involved in glycolysis and indirectly inhibited expression of genes involved in the Krebs cycle in *P. polymyxa* WLY78.

## DISCUSSION

Fnr is a global transcriptional regulator that controls a lot of genes expression in response to the transition from aerobic to anaerobic conditions in many bacteria. Although Fnr is well known in *E. coli* and *B. subtilis*, the function of Fnr in *Paenibacillus*, especially in N_2_-fixing Paenibacillus, is not known. *P. polymyxa* WLY78 that fixes nitrogen in anaerobic or microaerobic conditions has four *fnr* genes. Here, the functions of the *fnr* genes of *P. polymyxa* WLY78 in nitrogen fixation and other metabolisms were investigated.

We found that like *P. polymyxa* WLY78, *P. polymyxa* M1, *P. polymyxa* E681 and *P. polymyxa* SC2 have four *fnr* genes and each of the four *fnr* genes exhibited more than 90% identity with its corresponding *fnr*, suggesting that these bacterial strains have a common *fnr* gene ancestor. Whereas, some *Paenibacillus* species and strains, such as *P. polymyxa* EBL06, *P. polymyxa* Sb3-1 and *P. jamilae* NS115 have only a *fnr* gene whose predicted product Fnr shows higher (34.78%) identity with Fnr7 than with Fnr1, Fnr3 and Fnr5, suggesting that Fnr7 is conserved in *Paenibacillus*. However, Fnr1 and Fnr3 of *P. polymyxa* WLY78 have high similarity with both *B. subtilis* Fnr and *B. cereus* Fnr in sequence and structure, suggesting that Fnr1 and Fnr3 of *P. polymyxa* WLY78, *B. subtilis* Fnr and *B. cereus* Fnr have a common *fnr* gene ancestor.

Deletion of *fnr1* and *fnr3* genes of *P. polymyxa* WLY78 resulted to about 50% decrease of both growth rate and nitrogenase activity under anaerobic condition. Deletion of *fnr5* and *fnr7* genes led to a slight decrease of both growth rate and nitrogenase activity under anaerobic condition. The data suggest that the *fnr1* and *fnr3* genes play important roles in growth and nitrogen fixation under anaerobic conditions. However, the growth rates and nitrogenase activities of the multiple deletion mutants, such as Δ*fnr17,* Δ*fnr57* and Δ*fnr357*, were higher than single deletion mutants. The data implied that there might be some interactions among the four Fur proteins. Recently, specific interaction between Fnr1 and Fnr3 of *H. seropedicae* has been determined by using two-hybrid assays (37). Fnr1 and Fnr3 of *H. seropedicae* directly regulate discrete groups of promoters (Groups I and II, respectively), while Fnr3–Fnr1 heterodimers regulate a third group (Group III) promoters (37). Whether heterodimer is formed between Fnr1 or Fnr3 with Fnr5 or Fnr7 of *P. polymyxa* WLY78 needs to be studied in the near future.

In this study, Fnr1 with His_6_-tags at its N-terminus and Fnr3 with His_6_-tags at its C-terminus were expressed and purified in *E. coli* under aerobic condition. Both of the purified recombinant protein solutions did not exhibit the characteristic brown color, suggesting that Fe_4_-S_4_ cluster was oxidized by O_2_. However, EMSA showed that His-tagged Fnr1 and His-tagged Fnr3 of *P. polymyxa* WLY78 could bind to the promoter regions with the Fnr-binding site (5’-TGTGA-N6-TCACA-3’). Binding to the specific promoters suggest that the aerobically purified Fnr1 and Fnr3 proteins of *P. polymyxa* WLY78 were active forms. Similar report was found that both *B. cereus* Fnr tagged with His at its C-terminus (Fnr_His_) and Fnr tagged with Strep at its N-terminus (_Strep_Fnr) were active when expressed and purified in *E. coli* under oxic conditions. *In vitro,* the aerobically purified *B. cereus* Fnr as a monomer bound to the promoter regions of *fnr* itself, *resDE, plcR* and the structural enterotoxin genes *hbl* and *nhe* (38). Unlike *B. cereus* Fnr*, B. subtilis* Fnr existed in an inactive state under aerobiosis, due to the [4Fe-4S]^2+^ cluster of FNR being converted by O_2_ to a [2Fe-2S]^2+^. The *B. subtilis* Fnr formed stable dimer under aerobic and anaerobic conditions independent of Fe-S cluster formation, but DNA binding of Fnr was dependent on the presence of intact [4Fe-4S]^2+^ cluster (11). As a member of CRP/FNR family transcription factors, Fnr should function as a dimer *in vivo*. It is known that many transcription factors as a dimer bind to its specific DNA sites and there are two pathways to form dimeric protein-DNA complexes. Dimer pathway implies that the protein can dimerize first and then associate with DNA, and monomer pathway means that two protein monomers bind DNA sequentially and form their dimerization interface while bound to DNA (39). It was proposed that *B. cereus* Fnr takes a sequential monomer-binding pathway to form a dimer. But *B. subtilis* Fnr as a homodimer binds to its specific DNA-binding site and activates transcription (11). Our results suggest that Fnr1 and Fnr3 of *P. polymyxa* WLY78 behaved as *B. cereus* Fnr did. Thus, we deduce that *in vivo* Fnr1 and Fnr3 proteins of *P. polymyxa* WLY78 may bind to the specific promoter region by a sequential monomer-binding pathway just as *B. cereus* Fnr did.

Genome-wide transcription analysis showed that 301 genes, including 202 genes and operons, were differentially expressed in Δ*fnr13* compared to *P. polymyxa* WLY78 (Table S2). Similar reports were found that *E. coli* Fnr controlling the synthesis of up to 125 genes (40). Of the 301 genes, 116 were markedly up-regulated, indicating that they were directly or indirectly repressed by Fnr, and 185 were significantly down-regulated, suggesting that they were directly or indirectly activated by Fnr. Notably, the 9 genes (*nifBHDKENXhesAnifV*) within the *nif* operon in *P. polymyxa* WLY78 were significantly down-regulated 6.51-7.47 Log_2_FC. The data were consistent with the decreased nitrogenase activity of Δ*fnr13* mutant. qRT-PCR also confirmed that the *nifH* transcription in Δ*fnr13* mutant was obviously reduced. However, there was no predicted Fnr-binding site in the promoter region of the *nif* operon and EMSA also showed that Fnr1 or Fnr3 did not bind to the promoter region of the *nif* genes. These results indicated that Fnr1 and Fnr3 indirectly activated the expression of *nif* gene operon under anaerobic conditions. It is known that GlnR, a global regulator of nitrogen metabolism is required for *nif* transcription under anaerobic and nitrogen-limited condition. However, *glnRglnA* operon was up-regulated in Δ*fnr13* mutant, suggesting that *glnR* expression was not in coordination with *nif* expression. Also, EMSA showed that there was no binding of Fnr1 or Fnr3 to the promoter region of *glnR* with an Fnr-binding site. We do not know whether Fnr5 or Fnr7 could bind to the promoter region of *glnR.* In contrast to our results, the combined deletions in both the *fnr1* and *fnr3* genes in *H. seropedicae* led to higher expression of *nifA*, *nifB* and *nifH*, which was probably as a consequence of their influence on respiratory activity in relation to oxygen availability (16). It was shown that Fnr was required for relief of NifL inhibition in *K. oxytoca* under anaerobic conditions (14).

Fe is an essential element for nitrogenase. Our study showed that 36 Fe transporter genes in Δ*fnr13* mutant were significantly down-regulated compared to wild type. But there were no Fnr-binding sites in the promoter regions of these genes, suggesting that Fnr1 and Fnr3 indirectly induced expression of genes involved in uptake of Fe. Of the 36 Fe transporter genes, only *feoAfeoB* were involved in Fe^2+^ uptake and the other 34 genes belonged to Fe^3+^ transport systems. We deduce that the two forms of Fe^2+^ and Fe^3+^ coexisted in the culture of *P. polymyxa* WLY78 grown in anaerobic condition at neutral pH and then both types of Fe^2+^ and Fe^3+^ uptake systems were induced by Fnr. But we do not know how Fnr indirectly induced expression of genes involved in uptake of Fe^2+^ and Fe^3+^.

Since nitrogenase is very sensitive to oxygen, nitrogen fixation under anaerobic or microaerobic conditions. We found that the *ndh* gene (NADH dehydrogenase) and the *sdhCAB* genes (succinate dehydrogenase) that are the major dehydrogenase genes in the respiratory chain in Δ*fnr13* mutant were up-regulated under anaerobic condition. Other dehydrogenase genes, such as *yutJ* (NADH dehydrogenase), *yugK* (Probable NADH-dependent butanol dehydrogenase)*, hcaD* (NAD(FAD)-dependent dehydrogenases) and *ldh* (L-lactate dehydrogenase), were up-regulated. On the contrary, anaerobic electron acceptor genes *narGHJI* (nitrate reductase, Nar), *nasABCD* (nitrite reductase, Nas) and *narK* (nitrate/nitrite transporter, NarK) were down-regulated in Δ*fnr13* mutant. Importantly, expression of *resDE* whose promoter has an Fnr-binding site was down-regulated under anaerobic condition. The direct regulation of *resDE* by Fur was also reported in *B. cereus.* The data indicated that Fnr1 and Fnr3 inhibited expression of the genes involved in aerobic respiratory chain, and activated expression of anaerobic electron acceptor genes. These results also suggest that Fnr1 and Fnr3 provided O_2_ protection and energy for nitrogen fixation under anaerobic condition.

Nitrogen fixation is a process in which electrons originating from low potential electron carriers, such as flavodoxin or ferredoxin molecules were transferred to molecular N_2_. In *K. oxytoca,* the electron was produced by pyruvate: flavodoxin oxidoreductase (encoded by *nifJ*) during tricarboxylic acid cycle (TCA) and then a flavodoxin (encoded by *nifF*) mediated electron transfer to the Fe protein of nitrogenase (34). At present, we do not know the specific electron transfer system for nitrogen fixation in *P. polymyxa* WLY78. According to our previous study, *P. polymyxa* has several genes encoding flavodoxin or ferredoxin or oxidoreductase may be involved in electron transfer to nitrogenase. Here, we showed that *fldA* (flavodoxin), *fldB* (flavodoxin), *flr* (flavoredoxin), *ydfE* (flavoprotein oxygenases), *porG porA* (pyruvate:ferredoxin oxidoreductase), *ywcH3* (flavin-dependent oxidoreductases) and *ywcH1* (flavin-dependent oxidoreductases) were not differentially expressed. But *fldA* (flavodoxin), *hydA* (Fe-Fe hydrogenase), *hemN1* and *hemN3* (coproporphyrinogen-III oxidase) were down-regulated, suggesting that these genes may play important role in transferring electron to nitrogenase.

Taken together, the copy numbers of the *fnr* gene vary among different *Paenibacillus* species and different *P. polymyxa* strains*. P. polymyxa* WLY78 has four *fnr* genes encoding a global anaerobic regulator. The Fnr7 was conserved in different *Paenibacillus* species and strains. Fnr1 and Fnr3 of *P. polymyxa* WLY78 has more similarity to each other than to Fnr5 and Fnr7. Fnr1 and Fnr3 of *P. polymyxa* WLY78 also has high similarity with *B. subtilis* Fnr and *B. cereus* Fnr. EMSA showed that the aerobically purified Fnr1 and Fnr3 could bind to the specific target DNA *in vitro* as *B. cereus* Fnr did. We deduce that *in vivo* Fnr1 and Fnr3 of *P. polymyxa* WLY78 may bind to the specific promoter region by a sequential monomer-binding pathway to form a complex of a dimeric protein and DNA. Deletion of *fnr1* and *fnr3* led to a significant decrease of nitrogenase activity under anaerobic condition. Transcriptional analysis showed that Fnr1 and Fnr3 indirectly activate expression of the *nif* gene and Fe transported genes under anaerobic condition. Fnr1 and Fnr3 inhibit expression of the genes involved in aerobic respiratory chain and activate expression of genes responsible for anaerobic electron acceptor genes, which might provide O_2_ protection and energy for nitrogenase. In addition to Fnr1 and Fnr3, the function of Fnr5 and Fnr7 need to be studied in the future. This study not only reveals the roles of *fnr* genes in nitrogen fixation and electron transport, but also will provide a clue to clarifying the regulatory mechanisms of Fnr in nitrogen fixation in response to O_2_.

## MATERIALS AND METHODS

### Strains and media

*P. polymyxa* WLY78 used here was isolated from rhizosphere of bamboo by our laboratory (41). *P. polymyxa* and Δ*fnr* mutants were routinely grown at 30°C in LB or LD medium (per liter contains: 5 g NaCl, 5 g yeast and 10 g tryptone) with shaking. When appropriate, antibiotics were added in the following concentrations: 12.5 mg/ml tetracycline, 5 mg/ml erythromycin and 100 mg/ml ampicillin for maintenance of plasmids.

Nitrogen-deficient media were used for assay of nitrogenase activity. Nitrogen-deficient medium contained (per liter) 10.4 g Na_2_HPO_4_, 3.4 g KH_2_PO_4_, 26 mg CaCl_2_• 2H_2_O, 30 mg MgSO_4_, 0.3 mg MnSO_4_, 36 mg Ferric citrate, 7.6 mg Na_2_MoO_4_ ·2H_2_O, 10 μg p-aminobenzoic acid, 5 μg biotin, 4 g glucose as carbon source and 2 mM glutamate as nitrogen source (41).

### Nitrogenase activity assays

For nitrogenase activity assays, *P. polymyxa* WLY78 and Δ*fnr* mutants were grown in 50 ml LD media (supplemented with antibiotics when necessary) in 250 ml test tubes shaken at 250 rpm for 16 h at 30 °C. The cultures were collected by centrifugation, washed three times with sterilized water and then resuspended in nitrogen-deficient medium containing 2 mM glutamate to a final OD_600_ of 0.3–0.5. Then, 3-5 ml of suspension was transferred to a 26 ml test tube which was sealed with rubber stopper. The headspace in the tube was then vacuumed and filled with argon gas (42). After C_2_H_2_ (10 % of the headspace volume) was injected into the test tubes, the cultures were incubated at 30°C and with shaking at 250 rpm. After incubating for 4-8 h, 100 μl of gas was withdrawn through the rubber stopper with a gas tight syringe and manually injected into the gas chromatograph (HP6890) to quantify ethylene production. All treatments were in three replicates and all the experiments were repeated three or more times.

### β-galactosidase assays

To confirm whether deletion of fnr genes affect *nif* gene transcription, *P*. *polymyxa* WLY78 and 12 *fnr* mutants were transformed with a recombinant plasmid carrying the *nif* promoter-*lacZ* fusion (P*nif-lacZ* fusion) (19). β-galactosidase activity was assayed according to the method described by Wang et al (19).

### Identification and sequence alignment of *P. polymyxa* Fnr proteins

The sequences of Fnr1, Fnr3, Fnr5 and Fnr7 from *P. polymyxa* WLY78 were aligned with that of the Fnr of *Bacillus subtilis* subsp. subtilis str. 168 (Ref seq: NP_391612.1) and *Bacillus cereus* F4430-73 (Ref seq: KMP55664.1) using Clustal W software. The conserved domains in the Fnr proteins were investigated by sequence searching to the Pfam database (http://pfam.sanger.ac.uk/). The secondary structure elements in the Fnr proteins were defined by ESPript 3.0 algorithm (43).

### Phylogenetic analysis

In the non redundant NCBI database, amino acid sequences were obtained by performing a BLASTP search. Multiple gene alignments were carried out with molecular evolutionary genetics analysis (MEGA) (44). The neighbor-joining trees were constructed and 1,000 bootstraps were done by using the MEGA 7.0.14 software.

### Construction of Δ*fnr* mutants

Here, 12 Δ*fnr* mutants, including single mutants Δ*fnr1*, Δ*fnr3*, Δ*fnr5* and Δ*fnr*7, double mutants Δ*fnr13*, Δ*fnr17*, Δ*fnr35*, Δ*fnr37* and Δ*fnr57*, triple mutants Δ*fnr137* and Δ*fnr357*, and quadruple mutants Δ*fnr1357*, were constructed. The unmarked, single and multiple *fnr* deletion mutants were constructed via homologous recombination using the suicide plasmid pRN5101 as described previously (19). The upstream and downstream fragments flanking the coding region of *fnr1*, *fnr3 fnr5* and *fnr7* were PCR amplified from the genomic DNA of *P*. *polymyxa* WLY78, respectively. The primers used for deletion mutagenesis were listed in Table S3. The upstream and downstream fragments of four *fnr* genes were then fused with *Bam*H I /*Hind*III digested vector pRN5101 in Gibson assembly master mix (New England Biolabs), generating the four recombinant plasmids. Then, each of these recombinant plasmids was transformed into *P*. *polymyxa* WLY78 as described by (19), and the single crossover transformants were selected for erythromycin resistance (Em^r^). Subsequently, marker-free deletion mutants (the double-crossover transformants) were selected from the initial Em^r^ transformants after several rounds of nonselective growth at 39°C. The marker-free deletion mutants were confirmed by PCR amplification and DNA sequencing analysis. The multiple *fnr* deletion mutants were constructed via the same method in the single *fnr* deletion mutant background.

### Expression and purification of Fnr1 and Fnr3 in *E. coli*

The coding regions of *fnr1* and *fnr3* were PCR amplified from the genomic DNA of *P*. *polymyxa* WLY78, respectively. These PCR products were cloned into pET-28b(+) (Novagen, USA) to construct tagged Fnr proteins with His-tag at the N-terminus of Fnr1 and C-terminus of Fnr3 respectively, and then transformed into *E. coli* BL21 (DE3). The recombinant *E*. *coli* strains were cultivated at 37°C in LB broth supplemented with 50 μg/ml kanamycin until midlog phase, when 0.2 mM IPTG was added and incubation continued at 16°C for 8 hours. Cells were collected and disrupted in the lysis buffer (50 mM NaH_2_PO_4_, 300 mM NaCl, 10 mM Imidazole) by sonication on ice. Recombinant proteins NHis_6_-Fnr1 and Fnr3-CHis_6_ in the supernatant were purified on Ni_2_-NTA resin (Qiagen, Germany) according to the manufacturer’s protocol. Fractions eluted with 250 mM imidazole were dialyzed into binding buffer (20 mM HEPES pH 7.6, 1mM EDTA, 10 mM (NH_4_)_2_SO_4_, 1 mM DTT, 0.2% Tween 20, 30 mM KCl) for electrophoretic mobility shift assays (EMSA). Primers used here were listed in Table S3.

### Electrophoretic mobility shift assays (EMSAs)

EMSAs were performed as described previously using a DIG Gel Shift Kit (2nd Generation, Roche, USA) (19). The promoter fragments of predicted target genes or operons were PCR amplified from the genomic DNA of *P*. *polymyxa* WLY78. The primers used here and DNA fragment sizes were listed in Table S2. The DNA fragments were labeled at the 3’ end with digoxigenin (DIG) using terminal transferase, and used as probes in EMSAs. Each binding reaction (20 μl) consisted of 0.3 nM labelled probe, 1 μg poly [d(A-T)] and various concentrations (0, 0.05, 0.2, 2, 6 μM) of purified His-tagged Fnr (apo-Fnr) in the binding buffer. Reaction mixtures were incubated for 30 min at 25°C, analyzed by electrophoresis using native 5% polyacrylamide gel run with 0.5×TBE as running buffer at 4°C, and electrophoretically transferred to a positively charged nylon membrane (GE healthcare, UK). Labelled DNAs were detected by chemiluminescence according to the manufacturer’s instructions, and recorded on X-ray film.

### Bacterial RNA extraction and transcriptomic analysis

*P. polymyxa* WLY78 WT and Δ*fnr13* mutant were grown in nitrogen-deficient medium under anaerobic condition in 250 ml test tubes shaken at 250 rpm for 8 h at 30°C. The cultures were quickly collected by centrifugation at 4℃ under anaerobic condition and stored in liquid nitrogen for further use. This experiment was repeated three times.

For bacterial RNA extraction, bacterial cultures at each experimental time point were harvested and rapidly frozen in liquid nitrogen. Total RNAs were extracted with RNAiso Plus (Takara, Japan) according to the manufacturer’s protocol. Removal of genomic DNA and synthesis of cDNA were performed using PrimeScript RT reagent Kit with gDNA Eraser (Takara, Japan). The concentration of purified RNA was quantified on a Nanodrop ND-1000 spectophotometer (NanoDrop Technologies, Thermo FisherScientific, USA).

Illumina Hiseq 4000 sequencing from the total RNA was completed in Novogene Bioinfomatics Technology Company (Beijing, China) following a default Illumina stranded RNA protocol. Differential expression analysis of two groups (two biological replicates per condition) was performed using the DESeq R package (1.18.0) (45). DESeq provides statistical routines for determining differential expression in digital gene expression data using a model based on the negative binomial distribution. The resulting P-values were adjusted using the Benjamini and Hochberg’s approach for controlling the false discovery rate. The differences of transcript level with an adjusted P-value <0.05 determined by DESeq were considered to be significant and the genes were assigned as differentially expressed genes (DEGs). The DEGs were annotated using KEGG (Kyoto Encyclopedia of Genes and Genomes) database (http://www.genome.jp/kegg/). Gene Ontology (GO) enrichment analysis of differentially expressed genes was implemented by the GOseq R package, in which gene length bias was corrected. All of the raw reads are archived at the NCBI Sequence Read Archive (SRA) database (accession number: PRJNA596607). Transcriptional analysis was performed in triplicate and the reproducibility of the biological repeats was high (a mean R^2^=0.929).

### qRT-PCR analysis

Transcription levels of genes among *P*. *polymyxa* WLY78 and *fnr* deletion mutants were compared by quantitative real-time RT-PCR (qRT-PCR) analysis. Primers used for qRT-PCR are listed in Table S3. qRT-PCR was performed on Applied Biosystems 7500 Real-Time System (Life Technologies, USA) and detected by the SYBRGreen detection system with the following program: 95°C for 15 min, 1 cycle; 95°C for 10 sand 65°C for 30 s, 40 cycles. The relative expression level was calculated using ΔΔCt method and 16S rRNA was set as internal control. Triplicate assays using RNAs extracted in three independent experiments were performed for each target gene.

## Acknowledgments

This work was supported by the China Natural National Science Foundation (Grant No. 31770083) and the Tianjin Innovative Experimental Project for Young researchers (Grant No. 2020001).

